# Development of Remdesivir as a Dry Powder for Inhalation by Thin Film Freezing

**DOI:** 10.1101/2020.07.26.222109

**Authors:** Sawittree Sahakijpijarn, Chaeho Moon, John J. Koleng, Dale J. Christensen, Robert O. Williams

## Abstract

Remdesivir exhibits in vitro activity against SARS-CoV-2 and was granted approval for Emergency Use. To maximize delivery to the lungs, we formulated remdesivir as a dry powder for inhalation using thin film freezing (TFF). TFF produces brittle matrix nanostructured aggregates that are sheared into respirable low-density microparticles upon aerosolization from a passive dry powder inhaler. In vitro aerodynamic testing demonstrated that drug loading and excipient type affected the aerosol performance of remdesivir. Remdesivir combined with optimal excipients exhibited desirable aerosol performance (up to 93.0% FPF; 0.82μm MMAD). Remdesivir was amorphous after the TFF process, which benefitted drug dissolution in simulated lung fluid. TFF remdesivir formulations are stable after one-month storage at 25 °C/60%RH. In vivo pharmacokinetic evaluation showed that TFF-remdesivir-leucine was poorly absorbed into systemic circulation while TFF-remdesivir-Captisol® demonstrated increased systemic uptake compared to leucine. Remdesivir was hydrolyzed to the nucleoside analog GS-441524 in lung, and levels of GS-441524 were greater in lung with the leucine formulation compared to Captisol®. In conclusion, TFF technology produces high potency remdesivir dry powder formulations for inhalation suitable to treat patients with COVID-19 on an outpatient basis and earlier in the disease course where effective antiviral therapy can reduce related morbidity and mortality.

## 1. Introduction

The coronavirus disease 2019 (COVID-19) is an ongoing worldwide pandemic. As of September 2020, laboratory-confirmed cases have been reported in 213 countries and territories with more than 30 million reported cases and close to 1 million reported deaths [1]. Although this disease is asymptomatic to mild in most people, in some cases, it can develop into pneumonia, acute respiratory distress syndrome (ARDS) and multi-organ dysfunction [2]. While effective treatments for COVID-19 are urgently needed, including therapeutics and vaccines, currently no drugs are approved to treat COVID-19. Only one drug, remdesivir that is administered by intravenous (IV) injection, has been granted emergency use authorization in adult and pediatric patients hospitalized with severe disease. Therefore, additional therapeutics and routes of administration are needed to treat this respiratory virus.

Remdesivir (GS-5734), an investigational broad-spectrum antiviral agent, was developed by Gilead Sciences Inc. [3]. Remdesivir is a monophosphoramide prodrug that is intracellularly metabolized into an adenosine analog (GS-441524) [4]. Both GS-441524 and remdesivir (GS-5734) are metabolized into the active nucleoside triphosphate (GS-443902) by the host [5]. The adenosine nucleoside analog can inhibit viral RNA polymerases, evade proofreading by viral exonucleases, and subsequently decreases viral RNA production [4]. Previous studies demonstrated that remdesivir has broad-spectrum activity against members of the filovirus (e.g., EBOV, MARV) [6], CoVs (e.g., SARS-CoV, MERS-CoV) [7, 8], and paramyxovirus (e.g., respiratory syncytial virus, Nipah virus, and Hendra virus) [9].

Remdesivir was initially developed for the treatment of EBOLA virus disease [3]. Recently, in vitro studies showed that remdesivir was effective against COVID-19 in the human airway epithelial cell [10]. Short term outcomes of remdesivir were reported for 53 patients infected with COVID-19 treated in over 20 hospitals [11]. It was reported that overall clinical improvement was observed in 36 of 53 patients (68%) receiving a 10-day regimen of IV remdesivir [11]. However, 32 patients (60%) in the study experienced at least one adverse event, and 12 patients (23%) reported serious adverse events [11]. The most common adverse events included diarrhea, rash, septic shock, acute kidney injury, renal impairment, and hypotension [11, 12]. Several studies are ongoing, including two clinical studies evaluating the efficacy and safety of remdesivir for the treatment of COVID-19 respiratory disease in humans [13, 14]. In one study, 308 patients with mild to moderate COVID-19 were given remdesivir as a 200 mg loading dose on day 1, then 100 mg IV once-daily for 9 days [13]. In another study, the same regimen was given to 452 patients with severe clinical manifestations of COVID-19 [14].

The feasibility to deliver remdesivir via different administration routes (e.g., intravenous (IV) injection, intramuscular (IM) injection and oral administration) has been investigated [3]. Unfortunately, remdesivir is not suitable for oral delivery since the drug is mostly metabolized and cleared by first-pass metabolism, resulting in poor hepatic stability and poor bioavailability [3]. The delivery of remdesivir by the IM route also faces the challenge of variable release from muscle and slow acting of the active compound in peripheral blood mononuclear cells [3]. Therefore, remdesivir is administrated by IV injection since this route allows wide distribution to most tissues including kidney, kidney medulla, and liver [3].

For IV administration, remdesivir was developed in two dosage forms, a concentrate solution intended for dilution and administration by infusion and a lyophilized powder for reconstitution and dilution intended for administration by infusion [3]. Since remdesivir is practically insoluble in water, both formulations contain sulfobutylether-beta-cyclodextrin (SBECD) as a solubility enhancer by complexation [3, 15-17]. Since the excipient, SBECD, is cleared by the kidneys, remdesivir is contradicted in patients with several renal impairment [3]. A pharmacokinetic study showed that both formulations are interchangeable based on similar plasma concentrations [3]. Lyophilized remdesivir powder is stable when stored below 30 °C and has a shelf-life of 51 months, while remdesivir concentrate solution for dilution requires the product to be stored at freezing storage conditions (−25 °C to −10 °C) for a shelf-life of 48 months [3].

Currently, remdesivir is administered by injection into a vein as an infusion to treat patients with COVID-19 only in hospitals. However, due to the pandemic, many patients are not able to be hospitalized, and they do not have access to injectable administration of remdesivir. To provide remdesivir for other patients besides those most severely ill, more convenient and accessible dosage forms for different routes of administration must be quickly developed and tested so that patients have more options to get treated.

Intriguing strategies for future improvement of remdesivir include the development of inhaled dry powder of remdesivir for direct administration to the primary site of infection, the lungs. The mechanism of coronavirus infection is mostly in the respiratory tract, especially in the deep lungs. Hence, the inhalation route must be immediately pursued as the most promising route of administration to maximize direct delivery to the target site without first-pass metabolism, boost local antiviral activity in the lungs, and limit the potential for systemic side effects [18]. In addition, the cost of the drug can be reduced, and the supplies of the drug can be maximized, thus treating more patients due to less dose required by inhalation as compared to injectable forms. The treatment cost can also be decreased when administered by inhalation, since patients may not need to visit hospitals as is required to administer the IV injectable dose. Therefore, more affordable and early stage treatment can be provided to patients with inhaled remdesivir.

Nebulization of the current IV formulation in a diluted form is a potential method of pulmonary administration; however, the drug is prone to degrade by hydrolysis in aqueous solution to form the nucleoside monophosphate, which has difficulty penetrating cell membranes, thereby minimizing the antiviral activity in the lung cells [19]. Another concern is the use of SBECD as an excipient in inhalation dosage forms. Although cyclodextrin is not an excipient in an approved FDA inhaled product [20], several reports demonstrated the use cyclodextrin in inhaled formulations. Tolman et al. [21, 22] previously demonstrated that aerosolized voriconazole nebulized solution that contained a diluted form of the commercially available Vfend®, an IV voriconazole dosage form containing SBECD, is capable of producing clinically relevant lung tissue and plasma concentrations of the drug in animals. However, in this form, voriconazole in lung tissue was not able to be detected 6-8 h after administration of a single inhaled dose in the animal study by Tolman et al. [21], while voriconazole dry powder lasted longer in the lungs (up to 24 h) in an animal study by Beinborn et al [23]. Therefore, a dry powder formulation for inhalation can be more favorable with a smaller number of daily doses.

The advantages of dry powders for inhalation over liquids administered by nebulizers are not limited only to improve the stability of the drug during inhalation, but also dry powder inhalers provide the ease of maintenance and having a short administration time [24]. The inconvenience of nebulized inhalation solutions includes the use of a large nebulizer device requiring electric power and water, user manipulation for cleaning and operating, and long nebulization times. Also, the cost per dose and initial cost of the nebulizer are higher than a DPI device [24]. Hence, dry powder inhalation is an ideal dosage form for the treatment of COVID-19 in an outpatient setting, which would minimize the risk of spreading the virus to healthcare professionals.

Although several techniques have been used to prepare inhalable powders, including mechanical milling and spray drying, the advantages of thin film freezing (TFF) over other techniques rely on the ability to produce aerosolizable particles composed of brittle matrix, nanostructured aggregates. These are high surface area powders that are ideally suited for dry powder inhalation. TFF employs ultra-rapid freezing (on the order of 100-1,000 K/sec) such that precipitation (either as a crystalline nanoaggregate or amorphous solid dispersion) and particle growth of the dissolved solute can be prevented [25]. Subsequently, nanostructured aggregates are formed as a low-density brittle matrix powder [26], which is efficiently sheared into respirable low-density microparticles by a dry powder inhaler upon inhalation by the patient [27]. Despite a large geometric diameter (>10 microns), low-density microparticles composed of these nanostructured aggregates can be delivered to the deep lung with optimal aerodynamic diameters of <3 to 4 microns [27]. Additionally, it has been reported that the engineered particles with a geometric diameter greater than 10 microns can extend drug retention time in the lungs due to the ability of the deposited particles to escape from macrophage phagocytosis [28]. According to a recent study by Longest et al., computational models demonstrated that nanoaggregates are favorable for higher drug absorption efficiency and dose uniformity in the lungs, compared to microparticles [29]. Furthermore, TFF can produce amorphous high surface area powders with submicron primary particles, which can enhance the dissolution rate and subsequently improve the bioavailability of poorly water-soluble drugs like remdesivir in the lungs [30].

This work aims to apply the TFF technology to develop remdesivir dry powder formulations for inhalation administered by a commercially available dry powder inhaler. We hypothesize that the ultra-rapid freezing rate of the TFF technology will produce low-density, high porosity brittle matrix powders of remdesivir, which are aerosolizable by the shear forces generated from the passive dry powder inhaler, and thus allow high doses of remdesivir to be administered to the lung.

## 2. Materials and Methods

### 2.1. Materials

Remdesivir for formulation was purchased from Medkoo Biosciences (Research Triangle Park, NC, USA). Remdesivir, GS-441524, and its heavy isotope internal standards were purchased from Alsachim (Illkirch-Graffenstaden, France). Lactose monohydrate, leucine, polysorbate 20, acetonitrile (HPLC grade), and trifluoracetic acid were purchased from Fisher Scientific (Pittsburgh, PA, USA). D-Mannitol was bought from Acros Organic (Fair lawn, NJ, USA). Dipalmitoylphosphotidylcholine (DPPC) was purchased from Avanti Polar Lipid, Inc. (Alabaster, Alabama, USA). Cholesterol, albumin, transferrin, ascorbic acid and Hanks’ Balanced Salt solution (HBBS) were purchased from Sigma-Aldrich (St.Louis, MO, USA). Sulfobutylether-beta-cyclodextrin (SBECD, Captisol®) was kindly provided by CyDex Pharmaceuticals, Inc. High-resistance Monodose RS00 dry powder inhalers were kindly provided by Plastiape S.p.A. (Osnago, Italy).

### 2.2 Preparation of dry powder for inhalation using thin film freezing

Remdesivir and excipients (i.e., Captisol®, mannitol, lactose and leucine) were dissolved in a mixture of either acetonitrile/water (50/50 v/v) or 1,4-dioxane/water (50/50 v/v) at the solid contents shown in Table 1. The solutions were stored in the refrigerator at 2-8°C before the thin film freezing process. The solution was passed through an 18-gauge syringe needle and dropped from a height of approximately 10 cm onto a rotating cryogenically cooled stainless-steel drum. The solutions were frozen at a drum surface temperature of −100 °C. The frozen samples were collected in a stainless-steel container filled with liquid nitrogen and then transferred into a −80 °C freezer before drying in a lyophilizer. The frozen samples were primary dried at −40 °C for 20 hours, and ramped to 25 °C over 20 hours, and then secondary dried at 25 °C for 20 hours. The vacuum pressure was set at 100 mTorr for the whole drying cycle.

**Table 1.** List of TFF remdesivir compositions and parameters used for their preparation.

| Formulation | Drug loading (% w/w) | Excipient | Solid content (% w/v) | Solvent (v/v) | Processing temperature (°C) |
| --- | --- | --- | --- | --- | --- |
| F1 | 5 | Captisol® | 1.00 | Acetonitrile/water (50/50) | -100 |
| F2 | 20 | Captisol® | 1.00 | Acetonitrile/water (50/50) | -100 |
| F3 | 20 | Mannitol | 1.00 | Acetonitrile/water (50/50) | -100 |
| F4 | 20 | Lactose | 1.00 | Acetonitrile/water (50/50) | -100 |
| F5 | 20 | Leucine | 0.75 | Acetonitrile/water (50/50) | -100 |
| F6 | 50 | Captisol® | 0.30 | Acetonitrile/water (50/50) | -100 |
| F7 | 50 | Mannitol | 0.30 | Acetonitrile/water (50/50) | -100 |
| F8 | 50 | Lactose | 0.30 | Acetonitrile/water (50/50) | -100 |
| F9 | 50 | Leucine | 0.30 | Acetonitrile/water (50/50) | -100 |
| F10 | 80 | Captisol® | 0.30 | Acetonitrile/water (50/50) | –100 |
| F11 | 80 | Mannitol | 0.30 | Acetonitrile/water (50/50) | –100 |
| F12 | 80 | Lactose | 0.30 | Acetonitrile/water (50/50) | –100 |
| F13 | 80 | Leucine | 0.30 | Acetonitrile/water (50/50) | –100 |
| F14 | 100 | - | 0.25 | Acetonitrile/water (50/50) | –100 |
| F15 | 100 | - | 0.50 | 1,4-dioxane/water (50/50) | –100 |
| F16 | 100 | - | 1.00 | 1,4-dioxane/water (50/50) | –100 |
| F17 | 80 | Captisol® | 1.00 | 1,4-dioxane/water (50/50) | –100 |
| F18 | 80 | Mannitol | 1.00 | 1,4-dioxane/water (50/50) | –100 |
| F19 | 80 | Lactose | 1.00 | 1,4-dioxane/water (50/50) | –100 |
| F20 | 80 | Leucine | 1.00 | 1,4-dioxane/water (50/50) | –100 |

### 2.3. Drug quantification (HPLC)

The content of remdesivir was determined through analysis with a Thermo Scientific™ Dionex™ UltiMate™ 3000 HPLC system (Thermo Scientific, Sunnyvale, CA, USA) at a wavelength of 246 nm. A Waters Xbridge C18 column (4.6 × 150 mm, 3.5 μm) (Milford, MA) was used at 30 °C and a flow rate of 0.8 mL/min. The isocratic method was performed for 4.5 minutes using a mobile phase of 50:50 (%v/v) water –acetonitrile containing 0.05% (v/v) TFA. The retention time of remdesivir was approximately ∼3.5 minutes. Dimethyl sulfloxide: acetonitrile: water (10:60:30, v/v) was used as diluent. All analyses exhibited linearity in the range tested of 0.2–250 μg/mL. All chromatography data were processed by Chromeleon Version 7.2.10 software (Thermo Scientific, Sunnyvale, CA, USA).

For quantity and quality analysis of stability samples of TFF-remdesivir powders, Agilent 1220 Infinity II LC system equipped with Agilent 1290 Infinity II evaporative light scattering detector (ELSD) was utilized. Waters XBridge BEH C18 column (4.6 x 150 mm, 3.5 μm) (Milford, MA) was used at 30 °C and a flow rate of 0.8 mL/min. The gradient method with a mobile phase ratio of 5 to 95 % acetonitrile in water with 0.1% trifluoroacetic acid was used for total running time of 20 minutes. 10 μL of each sample was injected. The stability samples were monitored with 246 nm UV and ELSD to detect possible degradants. For ELSD, evaporator and nebulizer temperatures were set at 60 °C, and nitrogen gas flow was 1.6 L/min. The chromatography data were processed by Chemstation version C.01.10.

### 2.4 In vitro aerosol performance

The aerodynamic properties of the TFF powder samples were determined using a Next Generation Pharmaceutical Impactor (NGI) (MSP Corp, Shoreview, MN) connected to a High Capacity Pump (model HCP5, Copley Scientific, Nottingham, UK) and a Critical Flow Controller (model TPK 2000, Copley Scientific, Nottingham, UK). The pre-separator was not used in this study. To avoid particle bounce, the NGI collection plates were coated with 1.5% w/v polysorbate 20 in 100% methanol and then let the plates dry before analysis. A high resistance Plastiape® RS00 inhaler (Plastiape S.p.A, Osnago, Italy) containing size #3 HPMC capsules (V-Caps® Plus, Lonza, Morristown, NJ) and 1-3 mg of powder was attached to a USP induction port by a molded silicon adapter, and the powder was dispersed to the NGI at the flow rate of 60 L/min for 4 s per each actuation. The deposited powders from the capsule, inhaler, adapter, induction port, stages 1-7, and the micro-orifice collector (MOC) were collected by diluting with a mixture of DMSO/water/acetonitrile (10:30:60 v/v). The remdesivir content in the deposited powders was determined using the HPLC method as described in Section 2.3.

Copley Inhaler Testing Data Analysis Software (CITDAS) Version 3.10 (Copley Scientific, Nottingham, UK) was used to calculate the mass median aerodynamic diameter (MMAD), the geometric standard deviation (GSD), the fine particle fraction (FPF), and the emitted fraction (EF). The EF was calculated as the total amount of remdesivir emitted from the device as a percentage of total amount of remdesivir collected. The FPF of recovered dose was calculated as the total amount of remdesivir collected with an aerodynamic diameter below 5 μm as a percentage of the total amount of remdesivir collected. The FPF of delivered dose was calculated as the total amount of remdesivir collected with an aerodynamic diameter below 5 μm as a percentage of the total amount of remdesivir deposited on the adapter, the induction port, stages 1-7 and MOC.

### 2.5 X-ray powder diffraction (XRPD)

The crystallinity of the powders was evaluated using a benchtop X-ray diffraction instrument (Rigaku Miniflex 600 II, Woodlands, TX, USA) equipped with primary monochromated radiation (Cu K radiation source, λ = 1.54056 Å). The instrument was operated at an accelerating voltage of 40 kV at 15 mA. Samples were loaded in the sample holder and scanned in continuous mode with a step size of 0.02° over a 2θ range of 5–40 ° at a scan speed of 2 °/min, and a dwell time of 2 s

### 2.6 Modulated differential scanning calorimetry (mDSC)

Thermal analysis of powder samples was conducted using a differential scanning calorimeter Model Q20 (TA Instruments, New Castle, DE) equipped with a refrigerated cooling system (RCS40, TA Instruments, New Castle, DE). Samples of 2-3 mg were weighed and loaded into a T-zero pan. The pan with a T-zero hermetic lid were crimped, and a hole was drilled in the lid before placing the pan in the sample holder. To evaluate the glass transition and glass forming ability type of remdesivir unprocessed powder, samples were heat at a heating ramp rate of 10 °C/min from 25 °C to 150 °C, then cooled down to −40 °C, and then heated again to 250 °C. To investigate crystallinity of the TFF formulations, samples were heated from 25 °C to 350 °C with a heating ramp rate of 5 °C/min. The scans were performed with a modulation period of 60 s and a modulated amplitude of 1 °C. Dry nitrogen gas at a flow rate of 50 mL/min was used to purge the DSC cell throughout the analysis. Data were processed by TA Instruments Trios V.5.1.1.46572 software.

### 2.7 Scanning electron microscopy (SEM)

Scanning electron microscopy (Zeiss Supra 40 C SEM, Carl Zeiss, Heidenheim an der Brenz, Germany) was used to identify surface particle morphology of the TFF remdesivir powder formulations. A small amount from each of the powder formulations was placed on a carbon tape. A sputter coater was used to coat all samples with 15 nm of 60/40 Pd/Pt before capturing the images.

### 2.8 Nuclear Magnetic Resonance (NMR)

^1^H-NMR spectra were obtained using a Varian® NMR 400 MHz Spectrometer (Agilent Inc., Palo Alto, CA, USA) at 25 °C and were used to confirm the purity of remdesivir in the TFF powders, and to study interactions between excipients and remdesivir of TFF powders. Formulations F7, F8, F9, F10, F14 and unprocessed L-leucine and remdesivir drug substance were dissolved in dimethyl sulfoxide-d6 (DMSO-d6). Solutions were then transferred to 5 mm NMR tubes for NMR data acquisition. Chemical shifts were referenced to a residual solvent, DMSO, at 2.50 ppm. The NMR spectrum peaks of remdesivir in the formulations produced by TFF were compared to those of remdesivir unprocessed powder as received from the vendor. Excipients peaks in TFF-remdesivir powders were also compared to those of unprocessed excipients.

### 2.9 Dissolution test

Simulated lung fluid (SLF) was used as a dissolution medium. SLF contained dipalmitoylphosphotidylcholine (DPPC), cholesterol, albumin, transferrin, ascorbic acid and Hank’s balanced salt solution (HBSS). The SLF was prepared by two steps including a preparation of liposomal dispersion and freeze drying. In the first step, a liposomal dispersion was prepared by mixing 2.12 mL of 25 mg/mL DPPC and 5 μL of 200 mg/mL cholesterol in chloroform. Then, the solution was dried using an evaporator until the dry film was obtained. The thin film of lipid was dispersed by adding 8 mL of 57 mg/mL bovine serum albumin in water, 1 mL of 15 mg/mL transferrin in water, and 88.5 μL of 10 mM ascorbate solution. The mixture was sonicated until it fully dispersed. The dispersion was adjusted to 10 mL by adding 1M HBSS). The dispersion was frozen at −20 °C for 6 hours, and primary dried in the lyophilizer at −40 °C for 30 hours and ramp to 25 °C for 12 hours and finally secondary dried at 25 °C for 20 hours. The lyophilized SLF was stored at 2-8 °C and reconstituted with purified water upon dissolution testing.

Dissolution testing of F10, F11, F12, F13, F14 and unprocessed crystalline remdesivir was performed under sink conditions in SLF. An NGI (MSP Corp, Shoreview, MN) connected to a High Capacity Pump (model HCP5, Copley Scientific, Nottingham, UK) and a Critical Flow Controller (model TPK 2000, Copley Scientific, Nottingham, UK) was used to load aerosolized powder. Three 24 mm diameter filter papers were placed on stage 4 on the NGI to capture aerosolized powder to be used in the dissolution test. The transwell systems were modified as described in Rohrschneider et al. The filter papers were loaded in the donor compartment of the modified transwell systems. Reconstituted SLF was heated up to 37 °C before loading 1.5 mL in the six-well plate. The experiment was initiated after placing the donor compartment in the plate and adding 0.1 mL of SLF onto the filter papers. The modified transwell systems was placed on an orbital shaker at 100 rpm and 37 °C. Samples of 0.1 mL were collected at 10, 20, 30, 45, 60, 90, 120, 180, 240, 360, and 540 mins. Samples were centrifuged and diluted with DMSO/acetonitrile (10/90) to separate proteins in SLF. Supernatants were collected and analyzed by HPLC as described in section 2.3. Removed volumes were refilled with fresh SLF to maintain the dissolution volume. At the last time point, the filter membranes were removed and washed with DMSO/acetonitrile (10/90) to analyze the amount of undissolved drug.

### 2.10 Stability study

TFF powder of F10, F12 and F13 was transferred to a borosilicate glass vial. The vials were dried in a lyophilizer at 25 °C and 100 mTorr for 6 h, then backfilled with nitrogen gas before stoppering inside the lyophilizer. The vials were sealed with aluminum caps before packing in an aluminum foil bag with a desiccant. All samples were stored at 25 °C/60%RH. After one month, samples were collected and analyzed to determine physical stability and chemical stability. Additionally, samples were analyzed to determine aerosol performance.

### 2.11 In vivo pharmacokinetic study

#### 2.11.1 Single-dose dry powder insufflation

In vivo pharmacokinetic study was conducted using Sprague–Dawley rats in compliance with the Institutional Animal Care and Use Committee (IACUC; Protocol Number AUP-2019-00253) guidelines at The University of Texas at Austin. Rats weighing 200 to 250 g (average weight of 214.3 g) were housed in a 12-h light/dark cycle with access to food and water ad libitum and were subjected to one week of acclimation time to the housing environment.

Intratracheal administration was carried out using the dry powder insufflator (DP-4M model, Penn-Century Inc., Philadelphia, PA) connected to the air pump (AP-1 model, Penn-Century Inc., Philadelphia, PA). TFF powder was passed through a No. 325 sieve (45 μm aperture) to break down large aggregates. TFF remdesivir F10 and F13 were given to 5 rats per group. A precisely weighed quantity of sieved TFF powder was introduced into the sample chamber of the insufflator device. The dose was targeted at 10 mg/kg. Each rat was placed on its back at a 45 ° angle, and its incisors were secured with a rubber band. A laryngoscope was used to visualize the trachea and insert the insufflator device into the trachea. The sieved TFF powder was actuated into the lung using 3-5 pumps (200 μL of air per pump). The mass of powder delivered was measured by weighing the chamber before and after dose actuations.

Following insufflation, whole blood was collected at each time point (5, 15, 30, 2, 4, 8, and 24 hours) by cardiac puncture into heparinized vials. The blood samples were centrifuged at 10,000 rpm for 3 mins to obtain plasma. Lung were harvested after collecting blood at the last time point. Plasma samples and lungs were frozen and stored at −20 °C until analysis.

#### 2.11.2 Analysis of remdesivir and its metabolites in plasma and lung tissue

Plasma samples were prepared by combining 100 μL of plasma with 100 μL of methanol that contained 100 ng/mL of the heavy labeled internal standards for remdesivir and GS-441524, followed by vortexing and centrifuging at 12,000 rpm for 15 minutes. The supernatant was placed in a 96 well plate for LC/MS/MS analysis.

Lung tissue samples were cut in small pieces, added into a 2 mL tube with 3.5 g of 2.3 mm zirconia/silica beads (BioSpec Producs, Bartlesville, OK), and homogenized at 4800 rpm for 20 seconds. After homogenization, 1000 μL of methanol, containing 100 ng/mL internal standards for remdesivir and GS-441524 was added to the tube, and the tube was vortexed and centrifuged at 12,000 rpm for 15 minutes. The supernatant was placed in a 96 well plate for LC/MS/MS analysis. Calibration standards were prepared for plasma and lung tissue in the same fashion, spiking remdesivir and GS-441524 standard solutions into the blank plasma and lung tissue to obtain matrix matched calibrations. The calibration range was 0.1 _ 1000 ng/ml for plasma and 50 _ 10,000 ng/ml for lung tissue. The calibration range was chosen to bracket sample levels measured.

Remdesivir and GS-441524 were separated on an Agilent Poroshell column (2.1 x 50 mm, 2.7 μm) (Agilent, Santa Clara, CA) using a gradient of 0 to 90.25% of acetonitrile with 0.025% trifluoroacetic acid in 5 minutes at a flow rate of 0.35 mL/min, and the column temperature of 40°C. 10 μL of each sample was injected to analyze with an Agilent 6470 triple quadrupole LC/MS/MS system (Agilent, Santa Clara, CA).

#### 2.11.3 Pharmacokinetic analysis

The concentration of remdesivir and GS-441524 in plasma versus time was plotted. The data are presented as mean ± standard deviation (SD). The plots were used to compare the pharmacokinetic behavior of F10 and F13. A one compartmental analysis was performed to determine key pharmacokinetic parameters using Microsoft Excel 2016.

### 2.12 Statistical analysis

The statistical significance of experimental results was conducted using Student’s t-test in JMP 10. Alpha level was set at 0.05.

## 3 Results

### 3.1 Physical properties of TFF remdesivir

The particle morphology of TFF remdesivir powder formulations containing different drug loading concentrations and different excipients was determined using SEM. All formulations exhibited a brittle matrix structure of highly porous particles (Figure 1). For acetonitrile/water solvent system, Captisol®- and leucine-based formulations containing the same drug loading showed relatively smaller nanostructure aggregates, compared to mannitol- and lactose-based formulations. Different cosolvent systems resulted in different brittle matrix powder morphologies as observed by SEM. At the same drug load, mixture prepared from 1,4-dioxane/water produced smaller nanostructure aggregates, compared to an equivalent mixture prepared from acetonitrile/water (Figure 1B).

**Figure 1.**
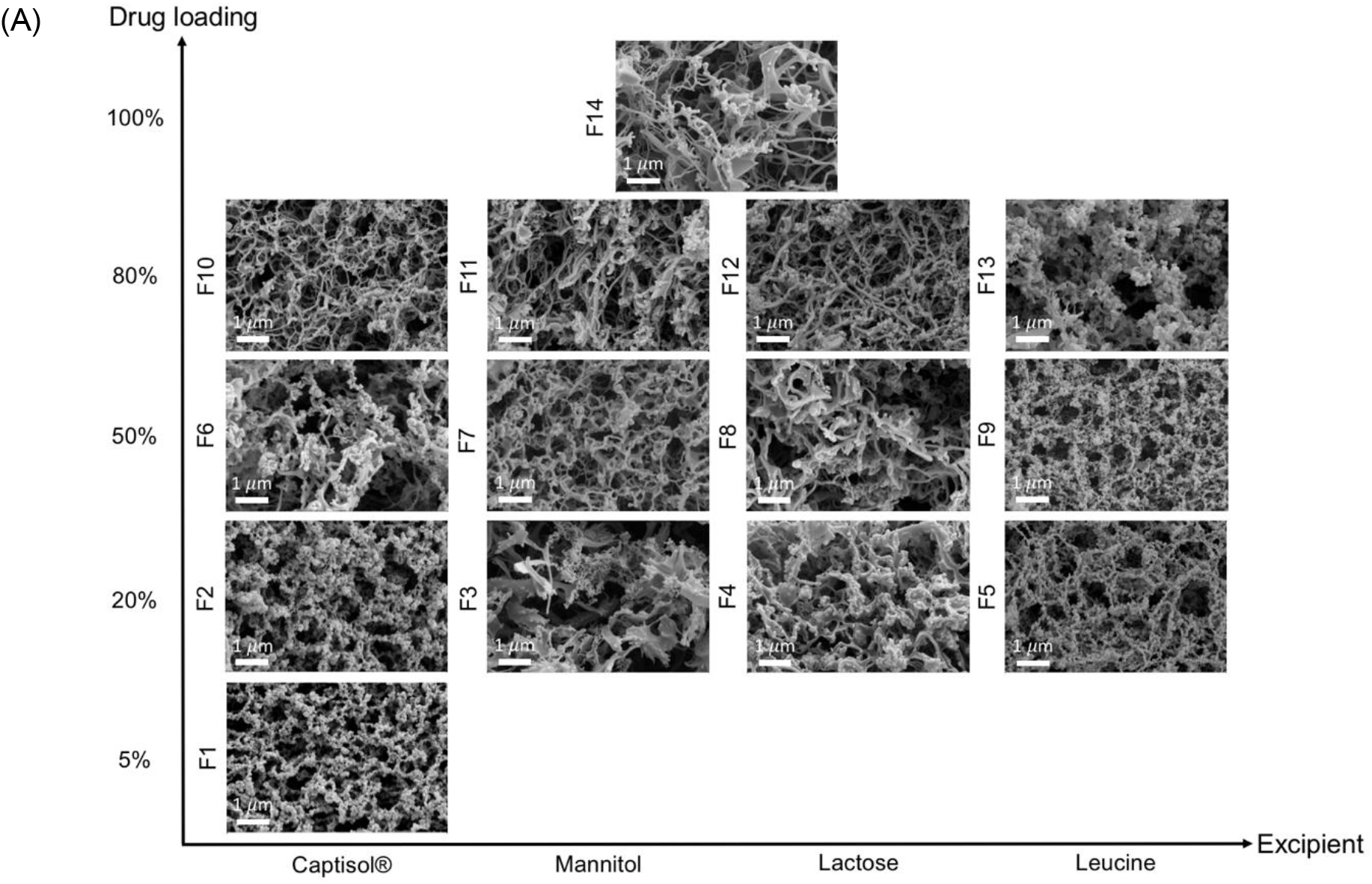
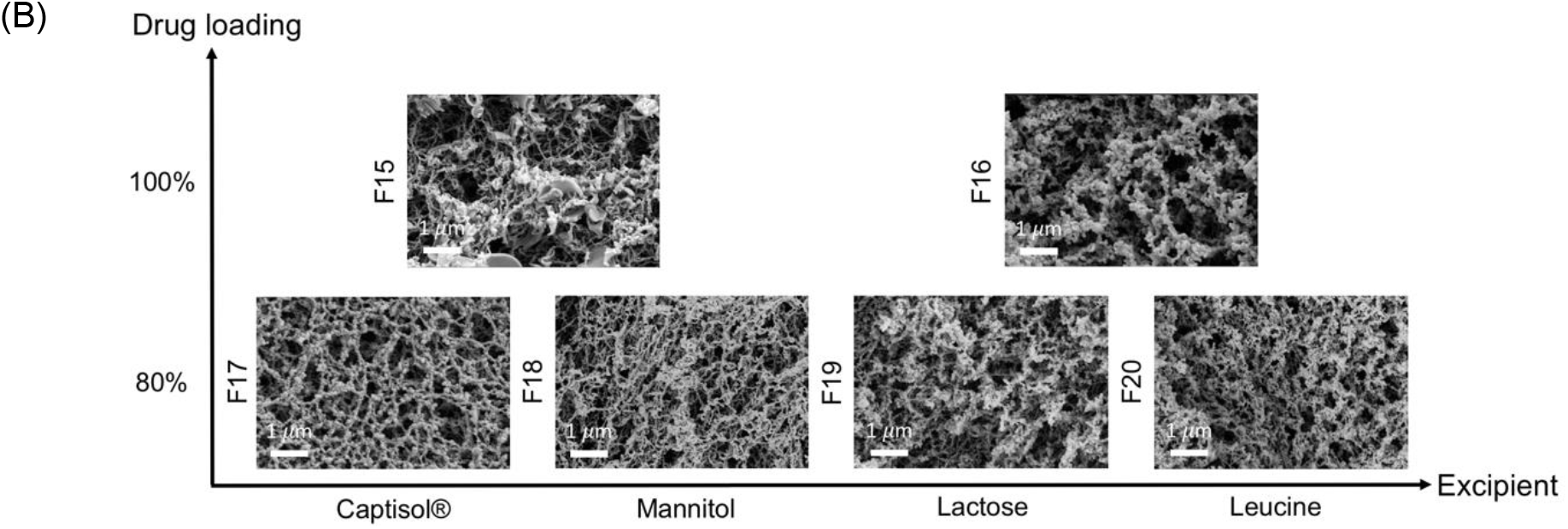
Brittle matrix particle morphology of TFF remdesivir compositions; (A) formulations prepared in acetonitrile/water (50/50) co-solvent mixture; (B) formulations prepared in 1,4-dioxane/water (50/50) co-solvent mixture.

Figure 2 shows the x-ray diffraction patterns of TFF remdesivir powder formulations. No sharp peaks of remdesivir were observed in any of formulations indicating remdesivir was amorphous after the TFF process. The drug loading and type of co-solvent did not affect the morphology of remdesivir. For excipients, sharp peaks of mannitol (13.5, 17, 18.5, 20.2,21, 22, 24.5, 25, 27.5, and 36 degree two-theta) were observed in F3, F7 and F11, indicating that mannitol in these formulations remained crystalline as a mixture of δ and α form [31]. Similarly, some peaks of leucine (6 and 19 degree two-theta) were observed in the TFF remdesivir – leucine formulations, F5, F9, F13, and F20, indicating that leucine remained crystalline after the process. In contrast, Captisol® and lactose in the compositions were amorphous after the TFF process.

**Figure 2.**
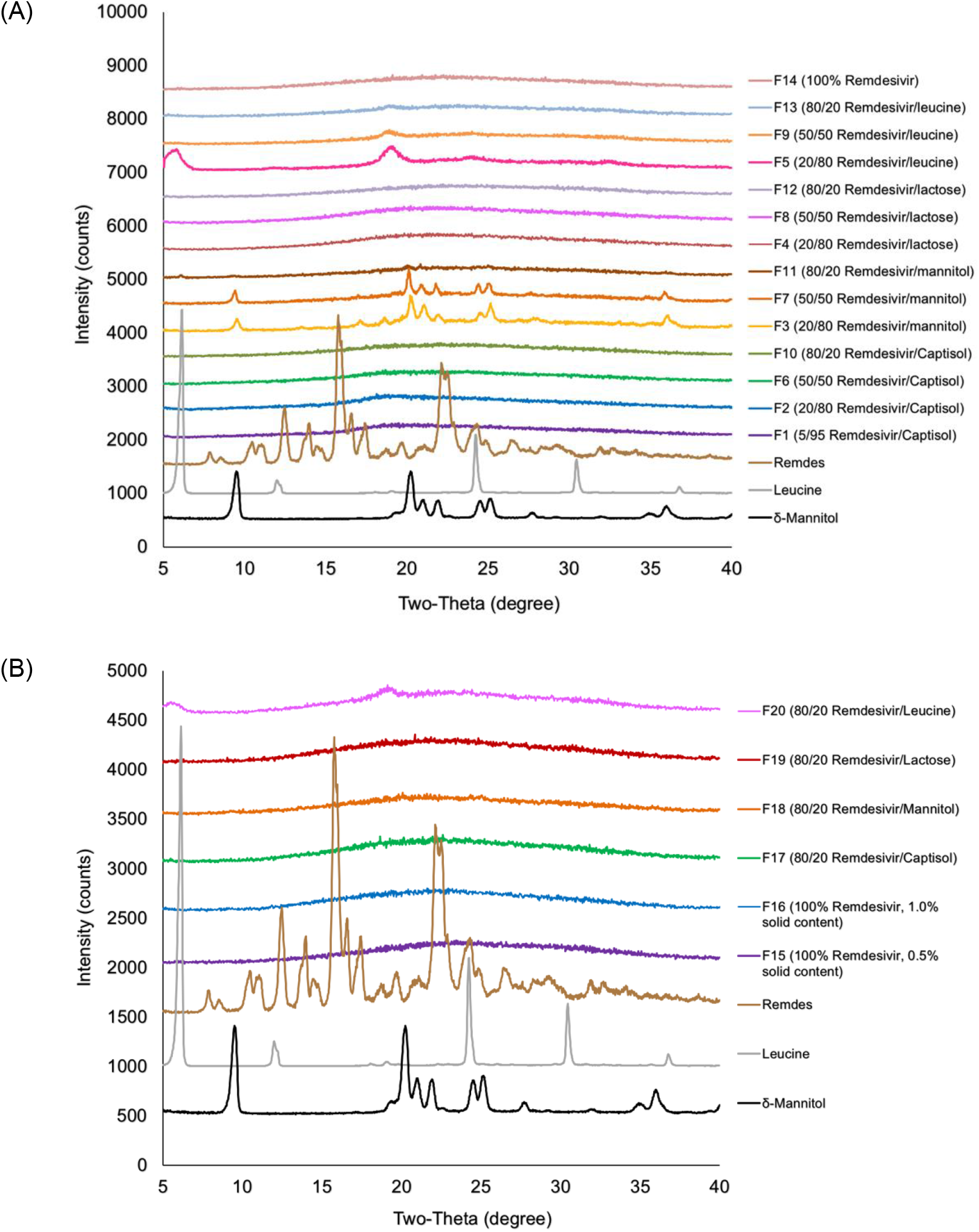
XRPD diffractograms of TFF remdesivir compositions; (A) formulations prepared in acetonitrile/water (50/50) solvent mixture; (B) formulations prepared in 1,4-dioxane/water (50/50) solvent mixture.

mDSC was employed to identify the glass-forming ability of remdesivir and determine the glass transition temperature (Tg) of remdesivir in each formulation. Figure 3A shows a mDSC thermogram of remdesivir unprocessed powder. The first heating cycle demonstrated remdesivir was crystalline with a melting point of ∼133 °C. The cooling and second heating cycle showed that the glass transition temperature of remdesivir was about 60 °C. No recrystallization peak was observed in any cycle.

**Figure 3.**
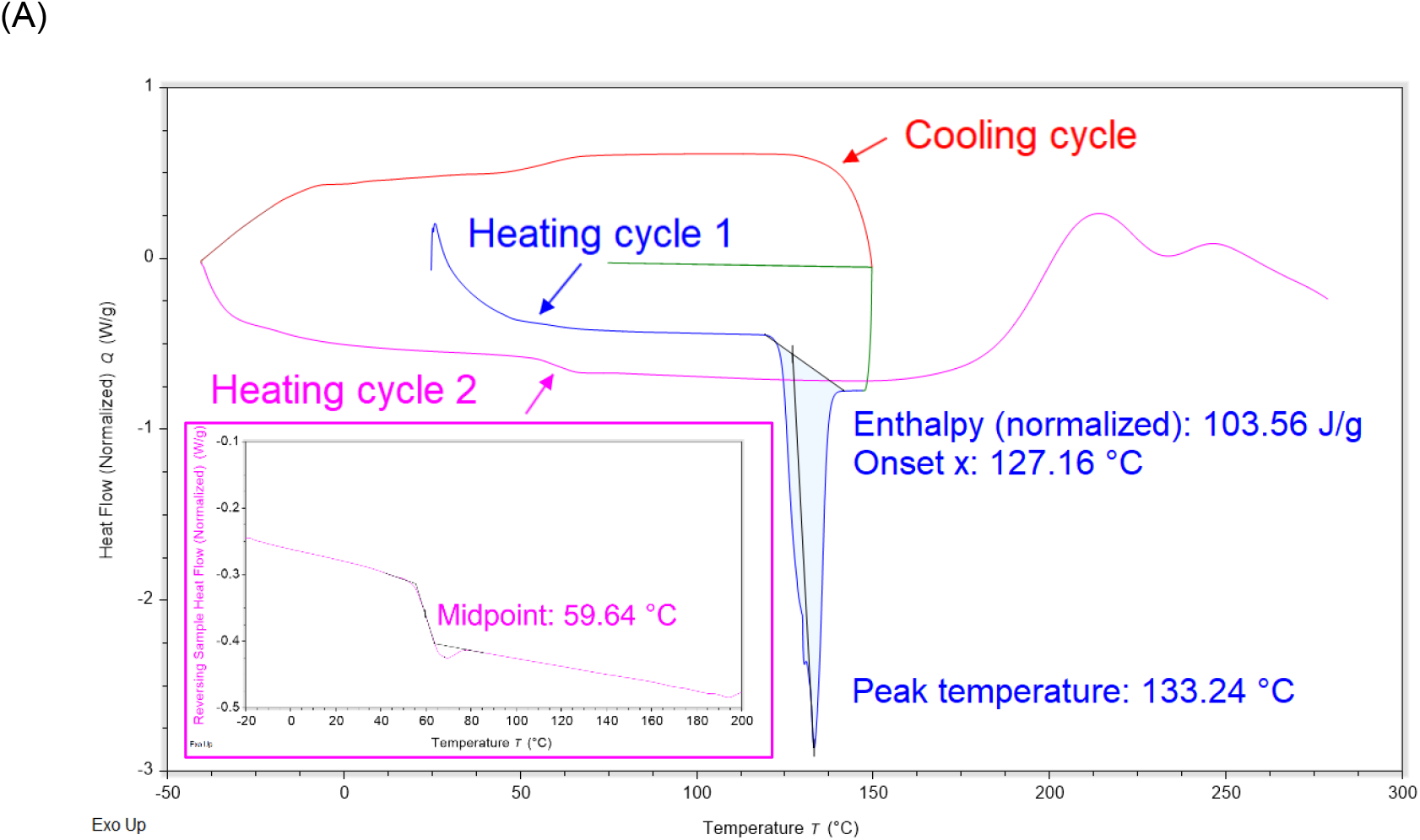
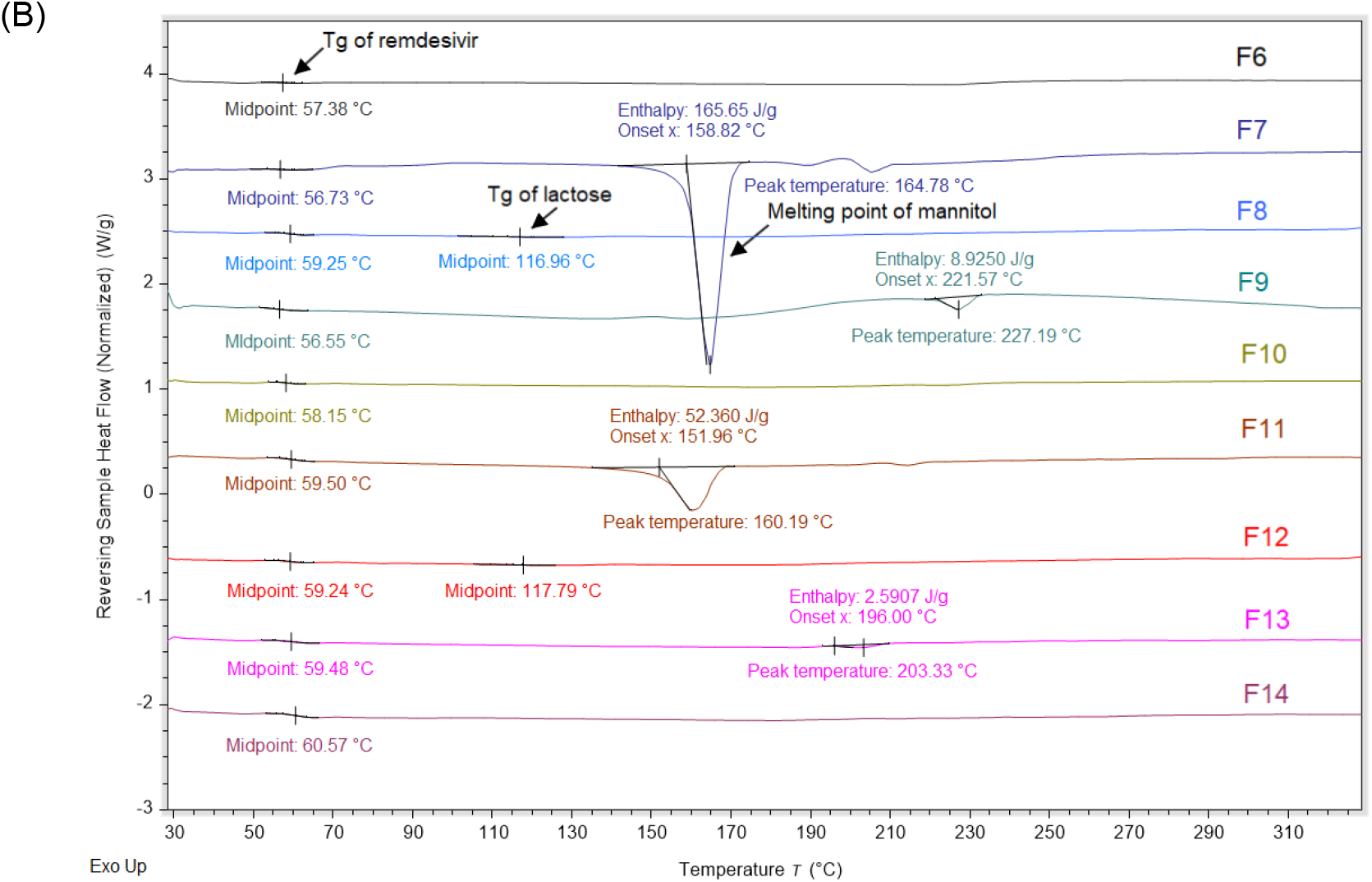
mDSC thermograms; (A) remdesivir unprocessed powder; (B) TFF remdesivir powder formulations.

Figure 3B shows the physical state of remdesivir and excipients. Despite the different formulation compositions, no melting peak of remdesivir was observed in any formulations, it remained amorphous. These results agree with XRD diffractograms showing that remdesivir was amorphous after the TFF process. The mDSC thermograms showed that the Tg of remdesivir in all formulations was in the range 56-61 °C. Captisol® in both F6 and F9 showed no endothermic peaks, demonstrating that Captisol® was in amorphous form. Endothermic peak of mannitol in F7 and F11 was observed at ∼165 °C and ∼160 °C, respectively, indicating mannitol was crystalline after TFF process. For lactose-based formulations, two glass transition temperatures were observed, including Tg of remdesivir (∼59 °C) and Tg of lactose (∼117 °C), indicating lactose and remdesivir were not molecularly miscible. For leucine-based formulations, an endothermic peak was observed in F9 and F13 at ∼227 °C and ∼203 °C, respectively, which are not identical with the melting peak of unprocessed leucine (∼300 °C) [32]. Therefore, the melting point depression of leucine can be attributed to an interaction of remdesivir and leucine.

### 3.2 Aerodynamic properties of TFF remdesivir

The aerodynamic particle size distribution of TFF remdesivir formulations was evaluated using a Plastiape® RS00 high resistance DPI and NGI apparatus. Figure 4 and Table 2 show aerodynamic diameter distribution and aerosol performance of TFF remdesivir formulations, respectively. In vitro aerodynamic testing revealed that drug loading affected the aerosol performance of the TFF formulations. Despite using different excipients, the aerosol performance generally increased as the drug loading was increased. The FPF (of recovered dose) of Captisol®-based formulations containing 5%, 20%, 50% and 80% of remdesivir (F1, F2, F6, F10, respectively) were 30.73 ± 1.11%, 45.84 ± 6.03%, 65.19 ± 1.18% and 68.29%, respectively. The MMAD of these TFF remdesivir powder formulations was 3.10 ± 0.04 μm, 2.59 ± 0.15 μm, 2.22 ± 0.14 μm, and 2.16 ± 0.21μm, respectively. For lactose- and mannitol-based formulations, there were significant increases in FPF (of recovered dose) when the drug loading was increased from 20% to 50%; however, FPF did not significantly change when the drug loading was increased further from 50% to 80% with these materials. Interestingly, FPFs of leucine-based formulations containing 20%, 50% and 80% of remdesivir were not significantly different. However, MMADs of these leucine-based formulations at drug loadings of 20%, 50% and 80% increased from 0.74 ± 0.06 μm to 0.82 ± 0.07 μm to 1.45 ± 0.07 μm, respectively. When the drug loading was increased to 100% (i.e., excipient free), F14 exhibited an FPF of 62.33% (of recovered drug) and an MMAD of 2.06 μm.

**Table 2.** Aerosol performance of TFF remdesivir powders using a Plastiape® RS00 high resistance DPI at a flow rate of 60 L/min (n=3). (MMAD, mass median aerodynamic diameter; GSD, geometric standard deviation; FPF, fine particle fraction; EF, emitted fraction)

| Formulations | MMAD<br>($\mu\text{m}$) | GSD | FPF<br>(%, recovered dose) | FPF<br>(%, delivered dose) | EF<br>(%, recovered dose) |
| --- | --- | --- | --- | --- | --- |
| F1 | $3.10 \pm 0.04$ | $2.88 \pm 0.03$ | $30.73 \pm 1.11$ | $36.87 \pm 0.78$ | $83.35 \pm 1.83$ |
| F2 | $2.59 \pm 0.15$ | $2.56 \pm 0.28$ | $45.84 \pm 6.03$ | $55.00 \pm 5.44$ | $83.32 \pm 6.70$ |
| F3 | $2.03 \pm 0.10$ | $2.76 \pm 0.07$ | $56.21 \pm 2.16$ | $62.27 \pm 2.72$ | $90.28 \pm 0.67$ |
| F4 | $2.02 \pm 0.37$ | $2.69 \pm 0.10$ | $52.38 \pm 4.65$ | $60.98 \pm 3.16$ | $85.86 \pm 5.40$ |
| F5 | $0.74 \pm 0.06$ | $3.50 \pm 0.13$ | $86.03 \pm 2.82$ | $92.10 \pm 1.76$ | $93.40 \pm 1.73$ |
| F6 | $2.22 \pm 0.14$ | $2.73 \pm 0.22$ | $65.19 \pm 1.18$ | $72.17 \pm 2.04$ | $90.37 \pm 2.49$ |
| F7 | $2.32 \pm 0.07$ | $2.57 \pm 0.03$ | $65.90 \pm 0.91$ | $73.08 \pm 0.22$ | $90.17 \pm 0.97$ |
| F8 | $2.40 \pm 0.39$ | $2.28 \pm 0.11$ | $68.43 \pm 4.33$ | $75.74 \pm 4.60$ | $90.34 \pm 0.82$ |
| F9 | $0.82 \pm 0.07$ | $3.19 \pm 0.10$ | $85.83 \pm 3.96$ | $92.99 \pm 1.11$ | $92.28 \pm 3.23$ |
| F10 | $2.16 \pm 0.21$ | $2.42 \pm 0.06$ | $71.48 \pm 5.52$ | $78.08 \pm 5.51$ | $91.48 \pm 2.09$ |
| F11 | $2.44 \pm 0.06$ | $2.53 \pm 0.06$ | $64.21 \pm 3.53$ | $71.62 \pm 3.10$ | $89.61 \pm 1.30$ |
| F12 | $2.03 \pm 0.11$ | $2.51 \pm 0.12$ | $70.32 \pm 3.39$ | $77.24 \pm 2.94$ | $91.02 \pm 1.55$ |
| F13 | $1.45 \pm 0.07$ | $2.17 \pm 0.01$ | $82.71 \pm 2.54$ | $89.68 \pm 0.91$ | $92.21 \pm 1.94$ |
| F14 | $2.09 \pm 0.07$ | $2.79 \pm 0.13$ | $65.80 \pm 2.68$ | $74.44 \pm 1.29$ | $88.37 \pm 2.16$ |
| F15 | $1.53 \pm 0.16$ | $2.70 \pm 0.10$ | $81.66 \pm 0.80$ | $85.11 \pm 1.20$ | $95.95 \pm 0.56$ |
| F16 | $1.42 \pm 0.20$ | $2.77 \pm 0.18$ | $84.25 \pm 1.87$ | $86.77 \pm 1.67$ | $97.10 \pm 0.40$ |
| F17 | $1.55 \pm 0.16$ | $2.99 \pm 0.43$ | $74.59 \pm 7.03$ | $78.76 \pm 6.75$ | $94.67 \pm 0.97$ |
| F18 | $2.03 \pm 0.20$ | $2.48 \pm 0.07$ | $75.34 \pm 1.06$ | $79.94 \pm 2.72$ | $94.30 \pm 1.94$ |
| F19 | $1.28 \pm 0.10$ | $2.92 \pm 0.17$ | $82.40 \pm 1.18$ | $87.28 \pm 0.57$ | $94.41 \pm 0.83$ |
| F20 | $1.29 \pm 0.11$ | $3.15 \pm 0.18$ | $82.26 \pm 3.86$ | $86.09 \pm 3.51$ | $95.53 \pm 0.60$ |

**Figure 4.**
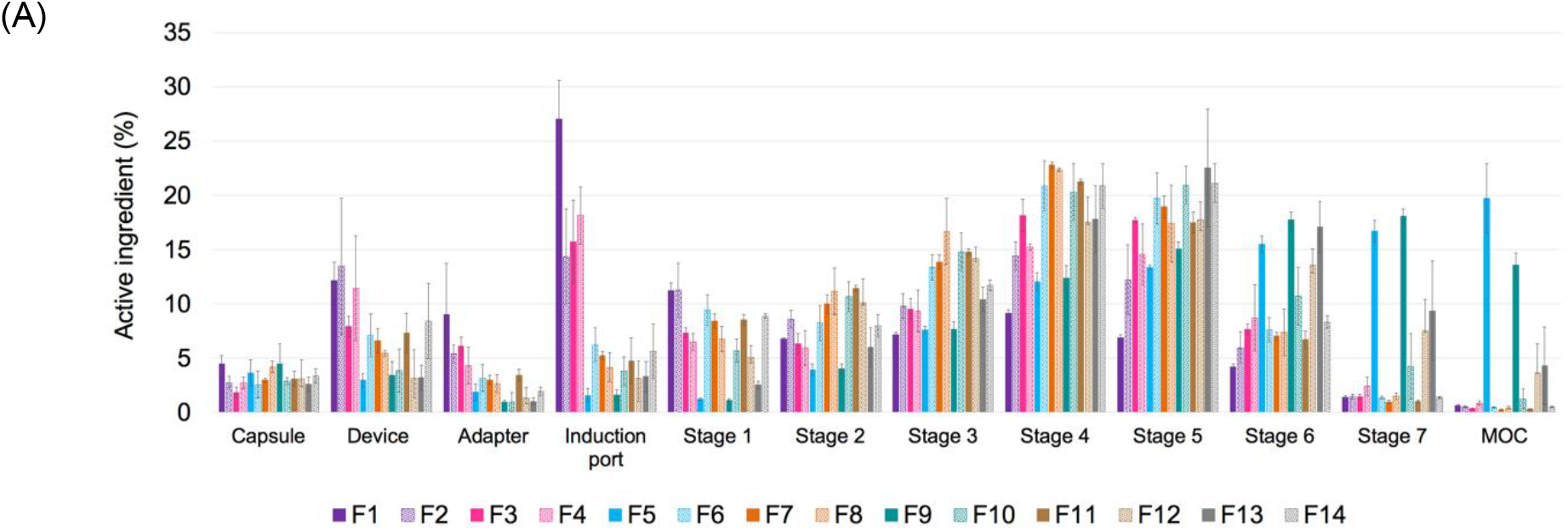
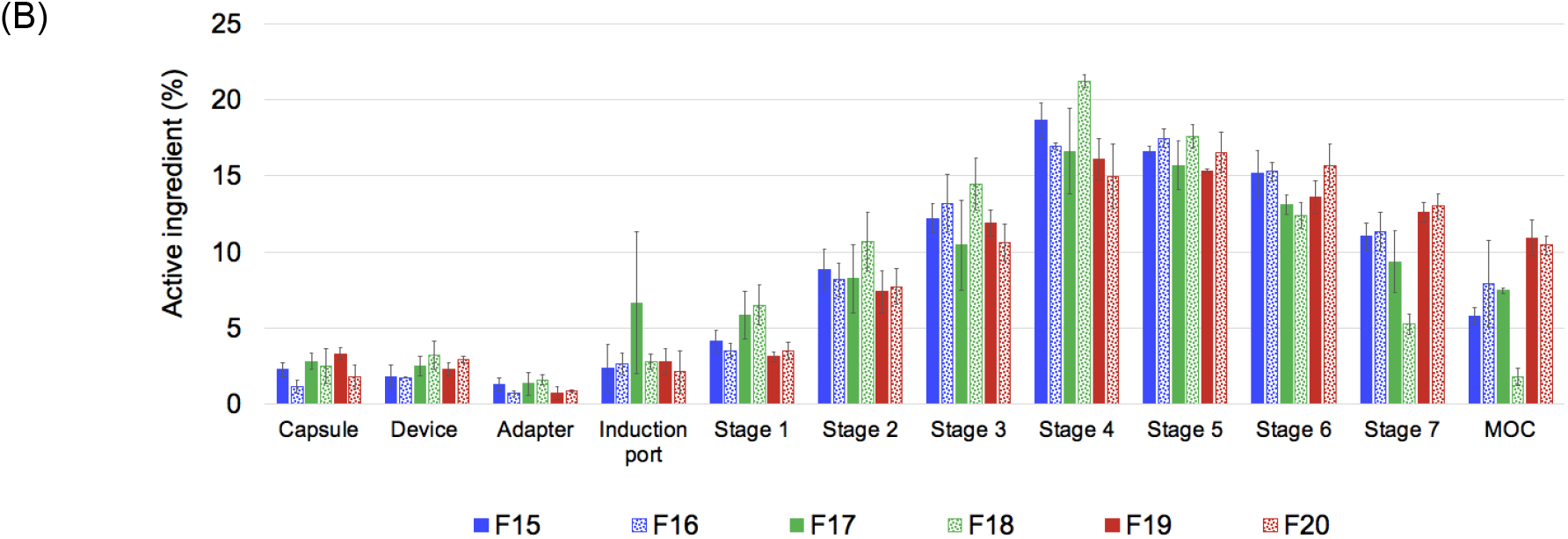
Aerodynamic diameter distribution of TFF remdesivir formulations when emitted from a Plastiape® RS00 high resistance DPI at a flow rate of 60 L/min (n=3); (A) formulations prepared in acetonitrile/water (50/50) co-solvent mixture; (B) formulations prepared in 1,4-dioxane/water (50/50) co-solvent mixture

An effect of the excipient on aerosol performance was also observed. At the same drug loading, leucine-based formulations exhibited significantly greater FPF (81-85% FPF), and smaller MMAD (0.7_1.5 μm), compared to other excipients, which indicates that leucine has distinct advantages over other compositions in terms of aerosol performance.

We also investigated the effect of co-solvent mixture and solid content on the aerosol performance of the formulations. For 100% drug loading (i.e., no excipient present in composition), significant differences in MMAD, FPF, and EF were observed between the formulations prepared from the acetonitrile/water (F14) and a 1,4-dioxane/water (F15 and F16) co-solvent systems. F15 and F16 exhibited smaller MMAD, larger FPF and EF, compared to F14. Comparing the formulations prepared in the same solvent system, F16 showed better aerosol performance although it was prepared at higher solid content.

Similar trends were observed in the Captisol®, mannitol, lactose, and leucine-based formulations. Captisol®, mannitol, lactose, and leucine-based formulations that were prepared in a 1,4-dioxane/water co-solvent (F17, F18, F19, F20, respectively) showed smaller MMAD, compared to the same compositions prepared in an acetonitrile/water cosolvent system (F10, F11, F12 and F13, respectively).

### 3.3 Interactions between TFF remdesivir and excipients

1H-NMR was performed to identify interactions between remdesivir and excipients. Figure 5B demonstrates an expansion of 1H-NMR spectra for selected TFF remdesivir powder formulations and remdesivir unprocessed powder. While the peak at 6.03 ppm is sharp and does not show any differences from the presented samples, the peaks at 5.37 and 6.33 ppm of F9 exhibited broader peak. Also, these peaks were slightly shifted to downfield. Figure 5C shows an expansion of 1H-NMR spectra of F9 and leucine unprocessed powder. The peak at 3.09 ppm of leucine was also shifted downfield to 3.13 ppm in F9. These results indicate that remdesivir and leucine form interactions during the TFF process.

**Figure 5.**
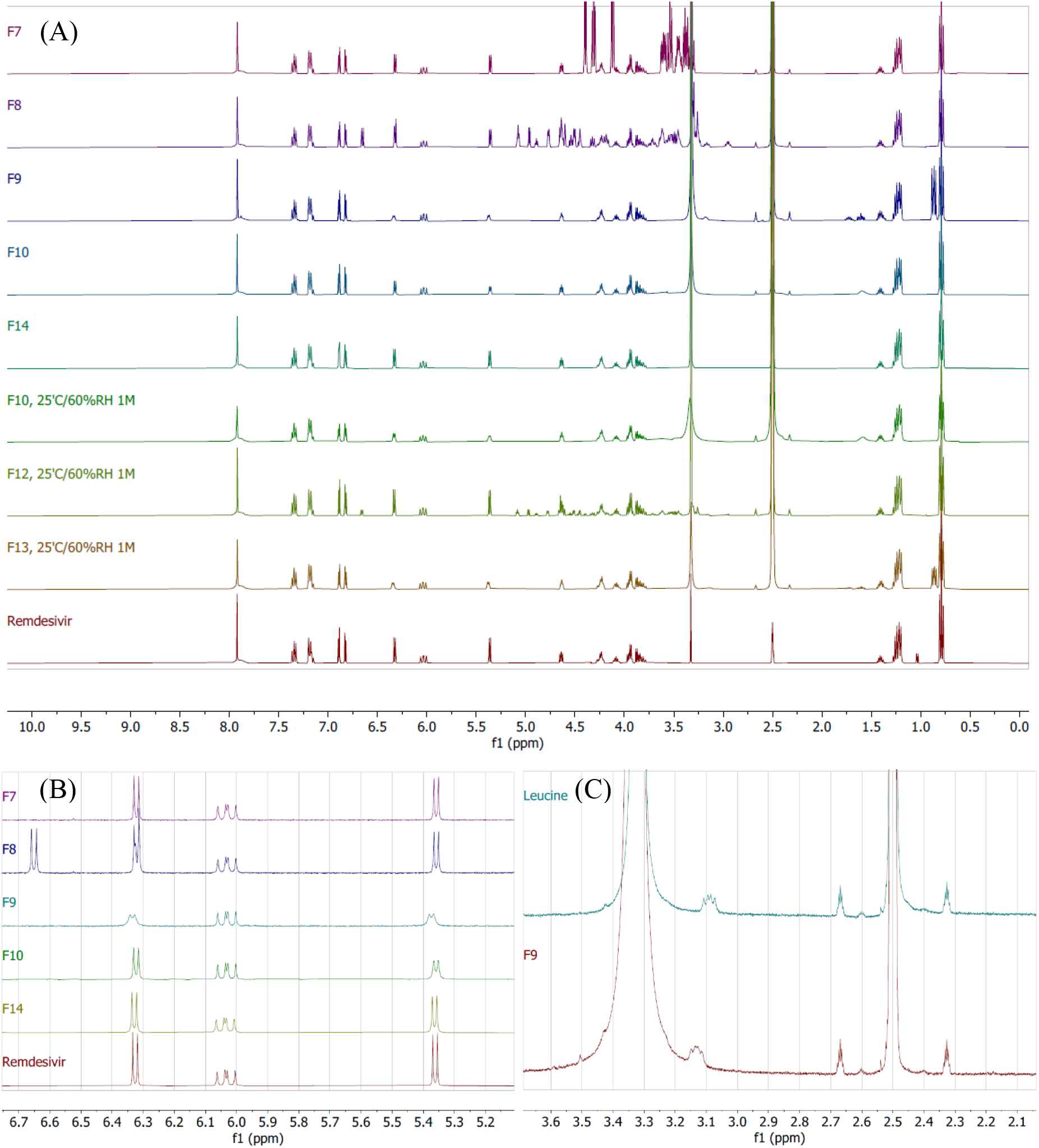
^1^H-NMR spectra; (A) TFF remdesivir powder formulations (F7, F8, F9, F10, F14) at initial condition, TFF remdesivir powder formulations (F10, F12, F13) at 25°C/60%RH at 1 month, and remdesivir unprocessed powder (Remdesivir); (B) expansion of TFF remdesivir powder formulations (F7, F8, F9, F10, F14) and remdesivir unprocessed powder (Remdesivir); (C) expansion of TFF remdesivir powder (F9) and leucine unprocessed powder (Leucine).

### 3.4 Stability study of TFF remdesivir

#### 3.4.1 Chemical stability of TFF remdesivir compositions

Chemical stability of remdesivir after TFF processing was confirmed by 1H-NMR. Figure 5A presents the 1H-NMR spectra of selected TFF remdesivir powder formulations and remdesivir unprocessed powder. Remdesivir peaks from the TFF remdesivir powder formulations consisting of 50% (w/w) remdesivir with mannitol, lactose, and leucine (F7, F8, and F9, respectively) are identical to the peaks of remdesivir unprocessed powder. The 80% (w/w) remdesivir with Captisol® (F10) and 100% remdesivir without excipient (F14) powders also presented indistinguishable peaks corresponding to remdesivir. Therefore, neither the aqueous-organic cosolvent mixture nor the conditions used in the TFF process affected the chemical stability of remdesivir.

After one-month storage at 25 °C/60% RH, samples were analyzed for chemical stability by NMR and HPLC. NMR spectra demonstrated that no chemical shift of remdesivir peaks and no new peaks were observed in F10, F12 and F13. Additionally, no degradant peak was observed in the HPLC chromatograms (data not shown), which agrees with NMR spectra. Both analyses indicated that remdesivir was chemically stable without degradation after storage.

#### 3.4.2 Physical stability of TFF remdesivir compositions

Samples after storage at 25°C/ 60%RH were analyzed for physical stability by XRD. XRD diffractogram demonstrates that no remdesivir peak was observed in any samples after storage (Figure 6), indicating that remdesivir in F10, F12 and F13 remained amorphous after one-month storage.

**Figure 6.**
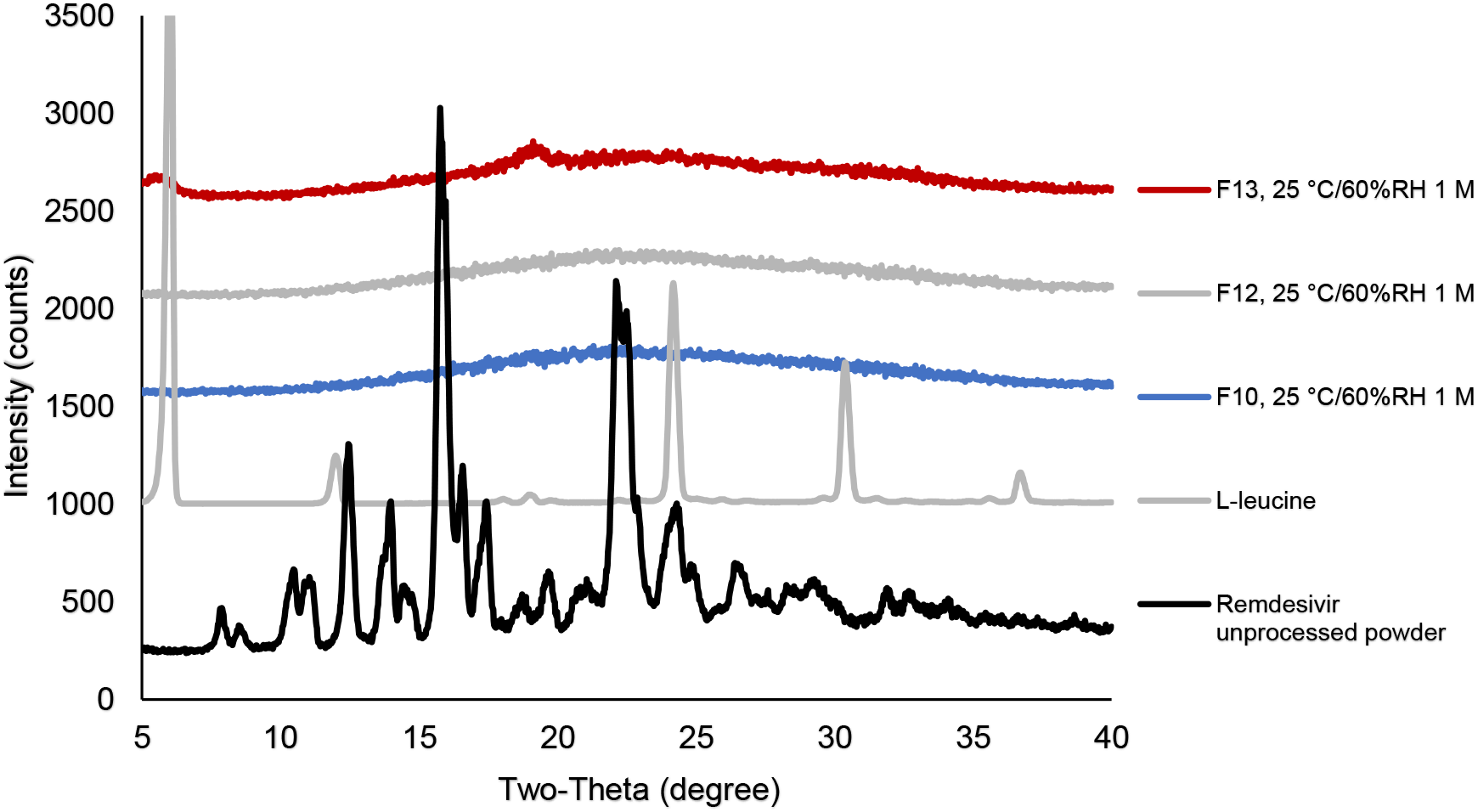
XRD diffractograms of F10, F12, and F13 after storage at 25 °C/60% RH.

#### 3.4.3 Aerosol performance after storage

The aerosol performance of F10, F12 and F13 was analyzed after one-month storage at 25 °C/60%RH. Overall, MMAD and FPF (of recovered dose) of F10, F12, and F13 was in the range of 1-2 μm and 74-83%, respectively (Table 3). Although MMAD was slightly decreased in all three formulations, there was no significant change in aerosol performance after one-month storage at 25 °C/60%RH.

**Table 3.** Aerosol performance of TFF remdesivir powders after one-month storage at 25 °C/60%RH using a Plastiape® RS00 high resistance DPI at a flow rate of 60 L/min (n=3). (MMAD, mass median aerodynamic diameter; GSD, geometric standard deviation; FPF, fine particle fraction; EF, emitted fraction)

| Formulation | Condition | MMAD<br>($\mu\text{m}$) | GSD | FPF<br>(%, recovered<br>dose) | FPF<br>(%, delivered<br>dose) | EF<br>(%, recovered<br>dose) |
| --- | --- | --- | --- | --- | --- | --- |
| F10 | Initial | $1.99 \pm 0.19$ | $2.49 \pm 0.13$ | $74.84 \pm 4.70$ | $80.22 \pm 3.08$ | $93.22 \pm 2.71$ |
| | 25 °C/60%RH,<br>1 M | $1.70 \pm 0.16$ | $2.67 \pm 0.13$ | $75.85 \pm 1.45$ | $80.40 \pm 1.79$ | $94.35 \pm 0.54$ |
| F12 | Initial | $1.78 \pm 0.23$ | $2.72 \pm 0.30$ | $76.88 \pm 6.37$ | $81.79 \pm 3.70$ | $93.88 \pm 3.61$ |
| | 25 °C/60%RH,<br>1 M | $1.41 \pm 0.20$ | $2.65 \pm 0.16$ | $81.46 \pm 2.32$ | $85.89 \pm 1.16$ | $94.84 \pm 1.52$ |
| F13 | Initial | $1.33 \pm 0.10$ | $2.54 \pm 0.42$ | $83.07 \pm 1.35$ | $88.19 \pm 2.43$ | $94.22 \pm 1.09$ |
| | 25 °C/60%RH,<br>1 M | $1.19 \pm 0.22$ | $2.96 \pm 0.16$ | $81.23 \pm 4.16$ | $87.60 \pm 1.82$ | $92.70 \pm 3.02$ |

### 3.5 Drug release in simulated lung fluid

The in vitro dissolution profile of remdesivir in SLF media under sink conditions are shown in Figure 8. The dissolution rates of TFF remdesivir formulations were significantly higher than the unprocessed crystalline remdesivir (p<0.05). More than 80% of remdesivir in F10, F11, F12 and F13 was released and dissolved within 90 mins, while 80% remdesivir in F14 was dissolved in 180 mins.

**Figure 7.**
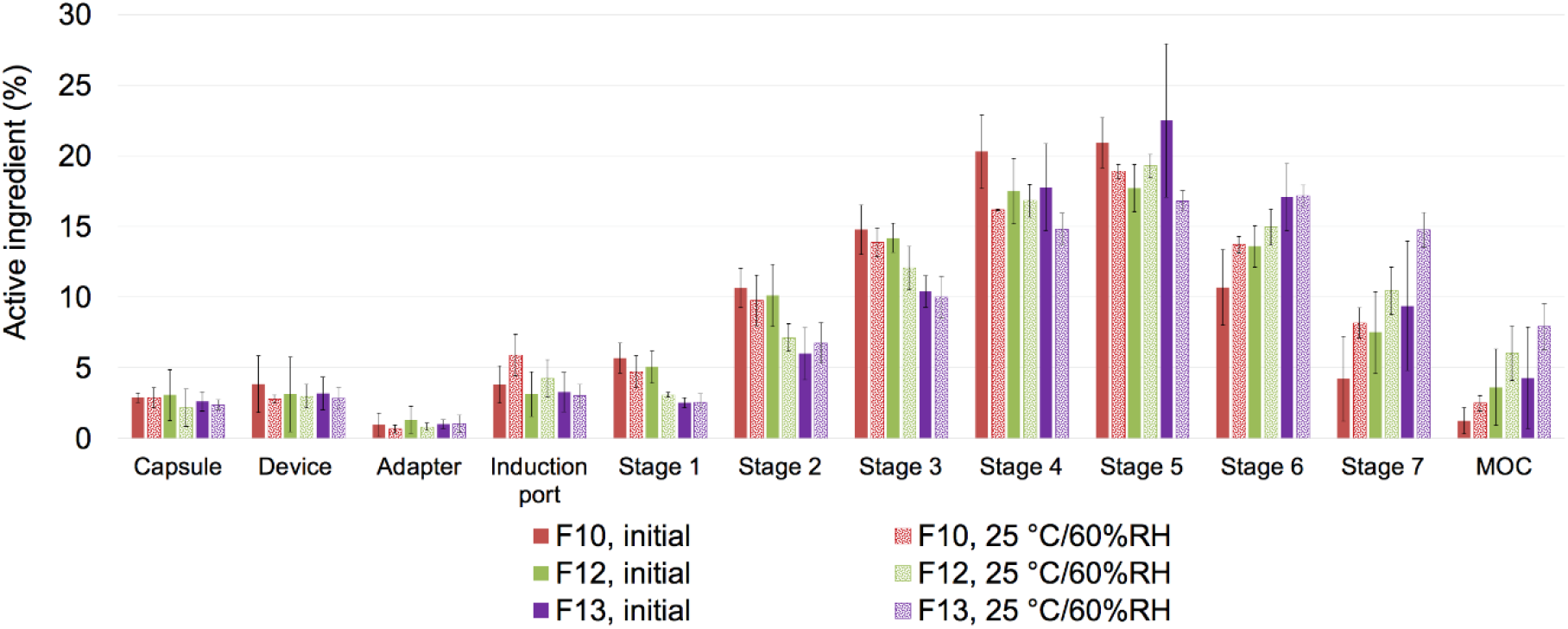
Aerosol performance of TFF remdesivir powder formulations after one-month storage at 25 °C/60%RH when emitted from a Plastiape® RS00 high resistance DPI at a flow rate of 60 L/min (n=3).

**Figure 8.**
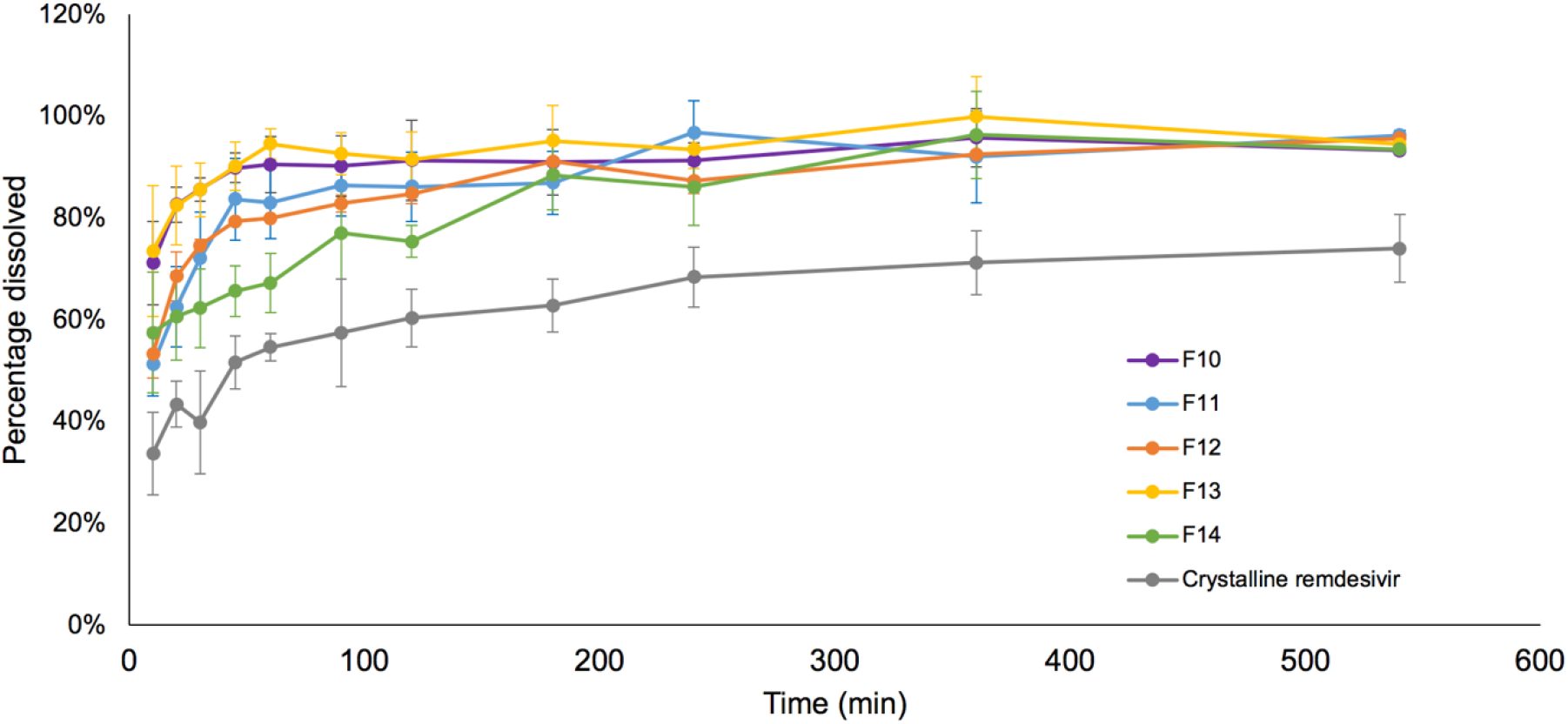
Dissolution profiles of TFF remdesivir dry powder compositions in simulated lung fluid.

### 3.6 *In vivo* pharmacokinetic study

The plasma concentration-time profiles of remdesivir and GS-441524 following a single inhalation dose of TFF remdesivir formulations are shown in Figures 9A and 9B, respectively. Remdesivir was detected in plasma at 5 min (0.068 ng/mL) and reached a maximum 30 min (1.149 ng/mL) following pulmonary dosing of F10. A much smaller amount of remdesivir (< 0.01 ng/mL) was detected at 5 and 15 mins following pulmonary dosing of F13. Remdesivir concentrations were below the lower limit of quantitation in plasma after 2 hours and after 30 minutes following pulmonary dosing of F10 and F13, respectively.

**Figure 9.**
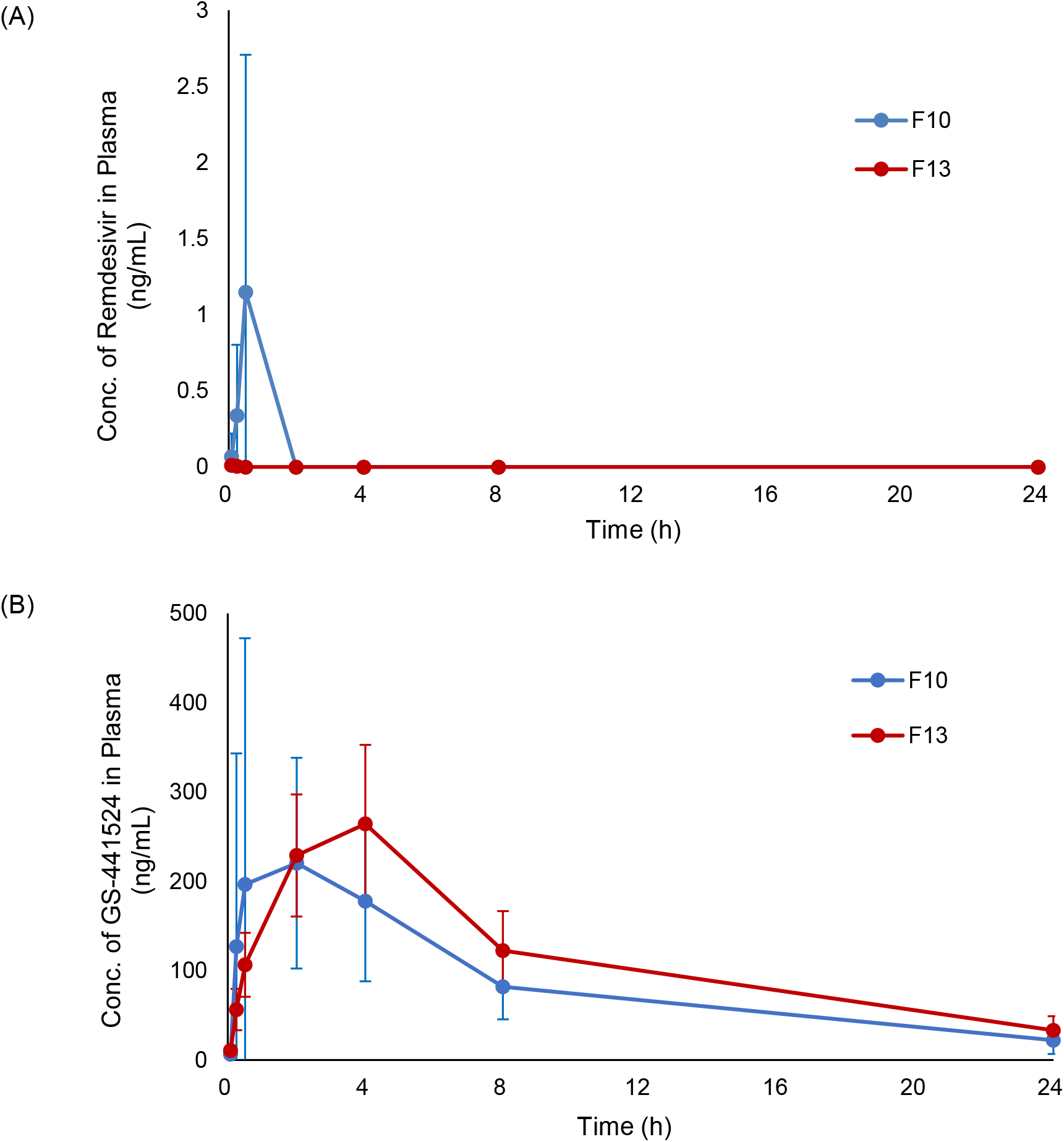
Plasma concentration - time profiles of F10 (Remdesivir-Captisol ®; 80/20 w/w) and F13 (Remdesivir-Leucine; 80/20 w/w) after a single inhalation administration in rats; (A) Remdesivir; (B) GS-441524.

Figure 9B shows a comparison of plasma GS-441524 concentration-time profile from F10 and F13, and the calculated pharmacokinetic parameters following pulmonary administration are presented in Table 4. The plasma GS-441524 profiles following pulmonary dosing of F10 and F13 has peak concentrations (C_max_) of 220.4 ± 118.0 ng/mL at 2 hours and 264.3 ± 88.5 ng/mL at 4 hours, respectively, before plasma concentrations started decreasing. AUC_0-24_ and AUC_inf_ of F13 were significantly greater than those of F10 (p<0.05).

**Table 4.** *In vivo* pharmacokinetic parameters for the plasma GS-441524 concentrations of F10 and F13 following a single inhalation administration.

| Formulations | T<sub>max</sub><br>(h) | C<sub>max</sub><br>(ng/mL) | AUC<sub>0-24</sub><br>(ng·h/mL) | AUC<sub>inf</sub><br>(ng·h/mL) |
| --- | --- | --- | --- | --- |
| F10 | 2 | 220.4 ± 118.0 | 2115.3 ± 979.6 | 2397.8 ± 1178.3 |
| F13 | 4 | 264.3 ± 88.5 | 2788.5 ± 857.1 | 3204.9 ± 967.6 |

**Table 5.** Amount of remdesivir and GS-441524 in the lungs at 24 hours following a single inhalation administration of F10 and F13.

| Formulation | Rat | Wet Lung<br>(mg) | Remdesivir |  | GS-441524 |  |
| --- | --- | --- | --- | --- | --- | --- |
|  |  |  | Total<br>Weight<br>(µg) | In wet Lung<br>(ng/mg) | Total<br>Weight<br>(µg) | In wet Lung<br>(ng/mg) |
| **F10** | A | 1164.5 | 16.5 | 14.2 | 6.2 | 5.3 |
|  | B | 1494.8 | 11.3 | 7.5 | 6.4 | 4.3 |
|  | C | 1255.2 | 13.6 | 10.9 | 10.8 | 8.6 |
|  | D | 1194.0 | 0.1 | 0.1 | 0.2 | 0.2 |
|  | E | 1176.3 | 16.2 | 13.8 | 12.2 | 10.4 |
|  | Average ± SD | - | 11.6 ± 6.7 | 9.3 ± 5.8 | 7.2 ± 4.7 | 5.8 ± 4.0 |
| **F13** | F | 1025.1 | 12.4 | 11.2 | 12.5 | 11.3 |
|  | G | 1126.8 | 7.7 | 7.6 | 15.3 | 14.9 |
|  | H | 1254.5 | 4.0 | 3.5 | 18.7 | 16.6 |
|  | I | 1263.1 | 3.4 | 2.7 | 18.7 | 14.9 |
|  | J | 1164.5 | 0.6 | 0.5 | 1.2 | 1.0 |
|  | Average ± SD | - | 5.6 ± 4.5 | 5.1 ± 4.3 | 13.4 ± 7.3 | 11.7 ± 6.3 |

Table 4 shows the amount of remdesivir and GS-441524 that remained in the lung tissues of treated animals at 24 hours following pulmonary administration of F10 and F13. Despite no statistical difference, F10 exhibited greater amounts of remdesivir in the lungs at 24 hours, while F13 showed greater amounts of GS-441524 in the lungs at 24 hours.

## 4. Discussion

### 4.1 Thin film freezing produces high potency remdesivir dry powders for inhalation with high aerosol performance

We investigated the feasibility of using the thin film freezing process to prepare inhalable remdesivir powder formulations. Different excipients including Captisol®, mannitol, lactose, and leucine were evaluated in this study. Mannitol and lactose are presently contained in FDA approved inhalation dosage forms, while leucine has gained recent interest for pulmonary delivery and is being used in an inhaled product in FDA clinical trials [33-35]. Captisol® was selected in this study since it is a solubilizer used in both the solution concentrate for dilution and infusion and the lyophilized powder for reconstitution/dilution and infusion of current remdesivir products [3].

The RS00 high resistance monodose DPI is a capsule-based DPI device that is available for commercial product development, and it functions to disperse the powder based on impaction force. A previous study confirmed that this impact-based DPI can disperse low-density brittle matrix powders made by TFF process into respirable particles better than a shear-based DPI (e.g., Handihaler®) [36]. Another study also evaluated the performance of different models of the monodose DPI (RS01 and RS00) on the aerosol performance of brittle matrix powders containing voriconazole nanoaggregates prepared by TFF [37]. It was shown that the RS00 device exhibited better powder shearing and deaggregation through smaller holes of the capsule created by the piercing system of the RS00 device [37]. Therefore, the RS00 high resistance Plastiape® DPI was selected for this study.

We found that excipient type and drug loading affect the aerosol performance of TFF remdesivir powder compositions. Overall, the aerosol performance of TFF remdesivir powders increased as the drug loading was increased, a highly desirable feature. This trend is obvious for the Captisol®-, lactose-, and mannitol-based formulations when the drug loading was increased from 20% to 50%. Furthermore, high potency TFF remdesivir powder without excipients (F14 and F15) also exhibited high FPF and small MMAD, which indicates remdesivir itself has a good dispersing ability without the need of a dispersing excipient when prepared using the TFF process. This shows that the TFF technology can be used to minimize the need of excipient(s) in the formulation, thus maximizing the amount of remdesivir being delivered to the lungs by dry powder inhalation.

The aerosol performance of the leucine-based formulations did not significantly change when the drug loading was increased from 20% to 80%, and these formulations exhibited superior aerosol performance compared to the other excipient-based formulations studied in this paper. This is likely attributed to the surface modifying properties of leucine. Several papers report that leucine can minimize the contact area and distance between particles [38, 39]. This decreases Van der Waal forces between drug particles and subsequently increases aerosol performance.

Additionally, different co-solvent systems affected the aerosol performance of TFF remdesivir powders. The formulations, except for the TFF remdesivir-leucine, prepared in a 1,4-dioxane/water co-solvent system exhibited smaller MMAD and higher FPF than the formulations prepared in an acetonitrile/water co-solvent system. Interestingly, the co-solvent systems did not affect the aerosol performance of the TFF remdesivir-leucine formulation. This agrees with SEM figures showing that the formulations prepared in a 1,4-dioxane/water co-solvent system has smaller nanostructured aggregates than the formulations prepared in an acetonitrile/water cosolvent system. This may be due to the difference in viscosity of the solvent system. The viscosity of ACN/water (50/50 v/v) was lower than that of 1,4-dioxane/water (50/50 v/v) (0.81 vs. 1.62 mPa·s) [40, 41]. Our results agree with previous studies by Beinborn et al. [42] and Moon et al. [37] showing that the viscosity of the solvent system has an impact on the aerosol performance of TFF powder. The higher viscosity of the co-solvent minimizes the movement of molecules during the ultra-rapid freezing step, resulting in more homogenous distribution in the frozen state [37]. On the contrary, the lower-viscosity solvent allows molecules to move more readily, which increases the chance of molecular aggregation and subsequently decreases the aerosol performance [37].

### 4.2 Physical and chemical stability of remdesivir dry powder produced using TFF

Both XRD diffractograms and DSC thermograms showed that remdesivir was amorphous after the TFF process. An amorphous form of the drug generally provides for faster dissolution rate than its crystalline form. Since the drug needs to be absorbed and then penetrate through the cell membrane before it is hydrolyzed to nucleoside monophosphate GS-441524, TFF remdesivir powders provide benefits for the dissolution of the deposited powder in the lung fluid that leads to improvement in the bioavailability and efficacy of the drug when administered by inhalation.

One potential concern related to the amorphous drug is physical instability due to its high energy state. According to criteria described by Wyttenbach et al., our study confirmed that remdesivir is categorized as a class III glass-forming drug (i.e., it is a stable glass former) [43], because no crystallization peak was observed in both the cooling and heating cycles on DSC. It was reported that the glass forming ability (GFA) has an influence on the physical stability of drug [44]. The GFA class III drugs have a strong tendency to transform into its amorphous state, resulting in the highest physical stability compared to class I and class II APIs [44, 45].

Additionally, NMR spectra demonstrated that Captisol® and leucine have interactions with remdesivir, which may help to stabilize remdesivir during storage. Also, DSC thermograms indicated that lactose and remdesivir were not molecularly dispersed in one single amorphous phase. To investigate an effect of drug and excipient interaction on physical and chemical stability, we selected F10, F12 and F13 in the stability study.

We found that the properties of a class III glass-forming drug agree with the physical stability of remdesivir. After one-month storage at 25°C/60% RH, the results showed remdesivir in all three formulations was physically stable. Different types of excipients did not affect the physical stability of remdesivir. Even though lactose was not molecularly dispersed with remdesivir, it’s use did not affect the physical stability of remdesivir. Lastly, these results indicate that amorphous remdesivir was physically stable without a stabilizer.

In addition to the physical stability, remdesivir, as a prodrug, is prone to degrade by hydrolysis in aqueous solution. Since an organic/water co-solvent system is required to dissolve the drug and excipients in the TFF process, chemical stability is another concern during preparation. NMR spectra demonstrated that remdesivir did not chemically degrade as a result of the TFF process. Even though remdesivir was exposed to binary co-solvent systems consisting of water during the process, the entire TFF process used to produce remdesivir dry powder inhalation formulations did not induce chemical degradation of the parent prodrug. Furthermore, remdesivir was chemically stable by HPLC (data not shown) and NMR after one-month storage at 25°C/60%RH.

### 4.3 Remdesivir in TFF powder compositions can be dissolved, absorbed and metabolized to GS-441524 in the lungs

Since remdesivir is a poorly water-soluble drug, its dissolution may be a critical factor of drug release in the lung fluid, especially in high drug load formulations. Undissolved particles can be cleared by mucociliary clearance or macrophage uptake, causing lower drug concentration, lung irritation, and inflammatory response [46]. Therefore, a dissolution test was evaluated in this study. The dissolution profile demonstrated that physical form of drug appears to have a significant effect on the dissolution rate. Amorphous remdesivir prepared using TFF had a much faster dissolution rate in simulated lung fluid as compared to crystalline remdesivir. Moreover, the high porosity and high surface area of the TFF powders also contribute to faster wetting, thereby enhancing the dissolution rate [30, 47].

The presence of excipients also affected the dissolution rate of remdesivir. The dissolution rate of 100% drug load formulation (i.e., no excipient was used), even though it is amorphous, was slower, compared to 80% drug loading formulations. The enhancement of dissolution rate is likely attributed to improved wetting from the use of excipients. Lactose and mannitol are hydrophilic carriers that can wet the powder formulations rapidly, promoting faster exposure of the drug particles to the dissolution medium [48]. In the case of Captisol®, cyclodextrins and their derivatives have been widely used as solubility and dissolution modifying excipients. In our study, the molar ratio of remdesivir to Captisol® in the high drug load composition (80/20 w/w) was 10:1, meaning that only about 10% of remdesivir was present in a complex with Captisol® in the solubilized form, the remaining remdesivir was not complexed with Captisol® and was present as amorphous powder in the matrix. It is known that certain drugs form a complex with Captisol® by hydrophobic interactions [15-17]. The inclusion complexation between cyclodextrin and drug can increase the aqueous solubility of drug, and increase the dissolution rate [49]. It was previously reported that the solubility of remdesivir can be increased by Captisol®, and Veklury®, the commercial product containing remdesivir, includes Captisol® to solubilize remdesivir for the IV solution formulation [15]. Interestingly, the TFF leucine-based formulation exhibited a higher dissolution rate than the TFF remdesivir formulations that contain mannitol or lactose, despite its hydrophobic properties. The faster dissolution rate of the leucine-based formulation is likely attributed to interactions (e.g., hydrogen bonding) between remdesivir and leucine. Peaks corresponding to acidic protons (-_OH or -_NH) of remdesivir from F9, a leucine-based formulation, became broader than those from the other formulations without leucine, and shifted downfield. These wider peaks result from inter-molecular interactions [50]. A peak shift in NMR spectra indicates magnetic field changes. The downfield shift of the acidic protons from F9 is caused by deshielded proton, reduced electron density, indicating hydrogen bonding at the protons [51, 52]. Along with a leucine corresponding peak shift from F9, it was concluded that remdesivir favorably interacts with leucine in hydrogen bonding. The molar ratio of remdesivir to leucine of the high drug load TFF compositions (80/20 w/w) was 1:1.5, meaning that leucine in this composition was in excess in terms of its ability to interact with remdesivir. Similarly, leucine interactions have been reported in Mangal’s study to enhance the solubility of azithromycin [53]. Specifically, leucine formed a complex with azithromycin through hydrogen bonding [53]. Generally, even though leucine is a hydrophobic amino acid, it is a water-soluble excipient that has higher solubility than poorly water-soluble drugs like remdesivir. Therefore, the interactions between drug and leucine at the molecular level can increase the dissolution rate of poorly water-soluble drug [53, 54], like that observed in our study.

Significantly higher plasma levels of GS-441524 than remdesivir were observed in our rat pharmacokinetic study. The half-life of remdesivir is reportedly much shorter than that of GS-441524. While half-life of GS-441524 is approximately 24.5 hours, for remdesivir it is only about 1 hour in humans following multiple once-daily IV administrations [3]. In rats, the half-life of remdesivir in plasma is reported to be less than 0.9 minute, which is much shorter than that in humans, due to higher esterase activity in rodent plasma [3]. Therefore, with a T_max_ of 30 minutes and a level of remdesivir in plasma that is lower than the detection limit at 2 hours, F10 presented an initial rapid systemic absorption of remdesivir that was complete at less than 2 hours. T_max_ of GS-441524 plasma concentration (2 hours) also supports this observation. In comparison, remdesivir plasma concentrations of F13 were lower than those of F10 at 5 and 15 minutes, and it was lower than the detection limit at the other time points, indicating that the initial systemic absorption of remdesivir from lung was much slower with F13. With the later T_max_ of GS-441524 plasma concentration, however, more sustained release of remdesivir from lung to plasma was observed.

The initial faster systemic absorption of F10 from the lungs is likely due to the fact that F10 contains about 10% remdesivir (on a molar basis) that is complexed with Captisol® in a solubilized form. Higher concentrations of both remdesivir and GS-441524 were detected in plasma following administration of F10, which contains 20% w/w of Captisol®. Captisol® (i.e., sulfobutylether-β-cyclodextrin), is known to enhance stability, solubility, and bioavailability of drugs [55]. When Captisol® forms a complex by hydrophobic interactions with drugs, these interactions are even greater than with simple physical mixtures of drug and Captisol® [16, 17]. Although F10 and F13 consist of amorphous remdesivir in the TFF remdesivir powder formulations, inclusion of Captisol® in F10 likely increased the absorption of the complexed remdesivir into systemic circulation due to its improvement in solubilization [15, 56]. Even though F10 incorporates a high molar ratio of drug to Captisol® (10:1), and not all of remdesivir in F10 complexes with Captisol®, partial remdesivir-Captisol® complex will still dissolve faster in lung fluid, rapidly becoming a solution and absorbed faster into systemic circulation than the non-complexed remdesivir fraction with Captisol® of F10. Eriksson et al. described that drugs in solution can be absorbed faster than dry powder in lungs in the isolated perfused lung model [57]. Enhanced absorption of inhaled drug complexed with Captisol® from lung to systemic circulation was reported in inhaled voriconazole studies. A solution form of nebulized voriconazole formulation with Captisol® resulted in an 8-fold higher AUC_plasma_ to AUC_lung_ ratio as compared to nanostructured amorphous voriconazole powder formulations when delivered to the lung directly (0.64 vs. 0.08 respectively) [21, 23].

Lastly, and related to potential use in COVID-19 therapy, the 50% maximal effective concentration (EC_50_) value is important to evaluate antiviral efficacy of drugs. While there are studies reported to evaluate remdesivir potency to inhibit SARS-CoV-2 replication, only one study has reported antiviral efficacy of GS-441524 against SARS-CoV-2. Pruijssers et al. determined inhibition of SARS-CoV-2 replication in established cell lines by GS-441524, and reported that EC_50_ is 0.62 μM (180.6 ng/mL) in Calu3 2B4 cells, and 0.47 μM (136.9 ng/mL) in Vero E6 cells [58]. The target dose of 10 mg/kg by dry powder insufflation in our rat pharmacokinetic study achieved these EC_50_ values necessary to reach EC_50_ to inhibit SARS-CoV-2. The plasma concentrations of GS-441524 in our pharmacokinetic study also achieved higher than reported EC_50_ of 0.18 μM (52.4 ng/mL) in SARS-CoV-infected HAE cells [59].

## 5. Conclusions

We have demonstrated that low density, highly porous brittle matrix particles of remdesivir powder formulations made by thin film freezing are highly aerosolizable and suitable for inhaled delivery to the lung, the site of SARS-CoV-2 replication. The properties of a good glass former, like remdesivir, supports that remdesivir maintains its physical stability during storage if protected from moisture. The thin film freezing process is suitable to prepare remdesivir dry powder for inhalation without causing chemical degradation. Dry powder administration can deliver remdesivir to the lungs, which appears to be converted to its nucleoside analog GS-441524 in the lung before entering system circulation based on the data showing negligible levels of remdesivir in the blood at all times for the TFF-remdesivir-leucine formulation. Therefore, dry powder remdesivir for inhalation produced using thin film freezing technology can provide a dry powder inhalation product for the treatment of patients with COVID-19 that can be administered on an outpatient basis and earlier in the disease course where effective antiviral therapy has a greater potential to reduce the morbidity and mortality associated with the disease.

